# Can charge-reversal be considered as a strategy for attaining thermal stability in proteins?

**DOI:** 10.1101/2021.05.09.443347

**Authors:** Suman Hait, Sudipto Basu, Sudip Kundu

## Abstract

Do charge reversal mutations (CRM) naturally occur in mesophilic-thermophilic/hyperthermophilic (M-T/HT) orthologous proteins? Do they contribute to thermal stability by altering charge-charge interactions? A careful investigation on 1550 M-T/HT orthologous protein pairs with remarkable structural and topological similarity extracts the role of buried and partially exposed CRMs in enhancing thermal stability. Our findings could assist in engineering thermo-stable variants of proteins.

**SIGNIFICANCE:** Protein engineering is one of the hot topics for decades specifically for its applications in different fields like de-novo protein design, directed evolution, making highly stable variants for food and drug industry etc. Proteins from organisms living in extreme environments are therefore a matter of common interest for scientists from different disciplines. Over three decades of study has already found several sequence and structural adaptations related to thermal stability, while charge reversal study remains ignored to a large extent. Influenced by nature’s strategy, our study provides a systemic understanding of how proper designing of few partially exposed and buried CRMs significantly contributes to thermal stability by altering the short distance electrostatic interactions.

## INTRODUCTION

Some proteins are considered useful in various research and industrial sectors because of their efficient and cost-effective catalytic properties. But naturally occurring proteins are not considered potential candidates for industrial applications because of their inadequate stability in various working conditions (1). Condition like extreme temperature demands special designing of proteins in thermophile and hyperthermophiles to function and survive. As proteins in thermophilic and hyperthermophilic organisms are more repellent to proteolysis and denaturation, engineering themostable variants of biocatalysts on the same mechanism that the nature uses (2–4), is a topic of immense interest for decades (5, 6). Among underlying molecular adaptations behind thermostability, number of charged amino acids is one of the few factors those exhibit a consistent incremental tendency from mesophilic to thermophilic and hyperthermophilic organisms (6). This motivates to look for the possible role of electrostatic interaction in thermostability. Previous studies show that mutations of amino acids in the positions involved in attractive (favourable for stability) electrostatic interactions can destabilize the structure (7), while removal of repulsive (unfavourable for stability) interactions can enhance stability (8). Computational 3D designing and rational optimization of charge-charge interactions for designing thermostable proteins have also been reported in several literature (9–13). Apart from these studies, the instances of charge reversal studies are rare, despite the fact that a single CRM can change local interaction pattern to a large extent (**Fig 1A,B**).

**Fig 1.**
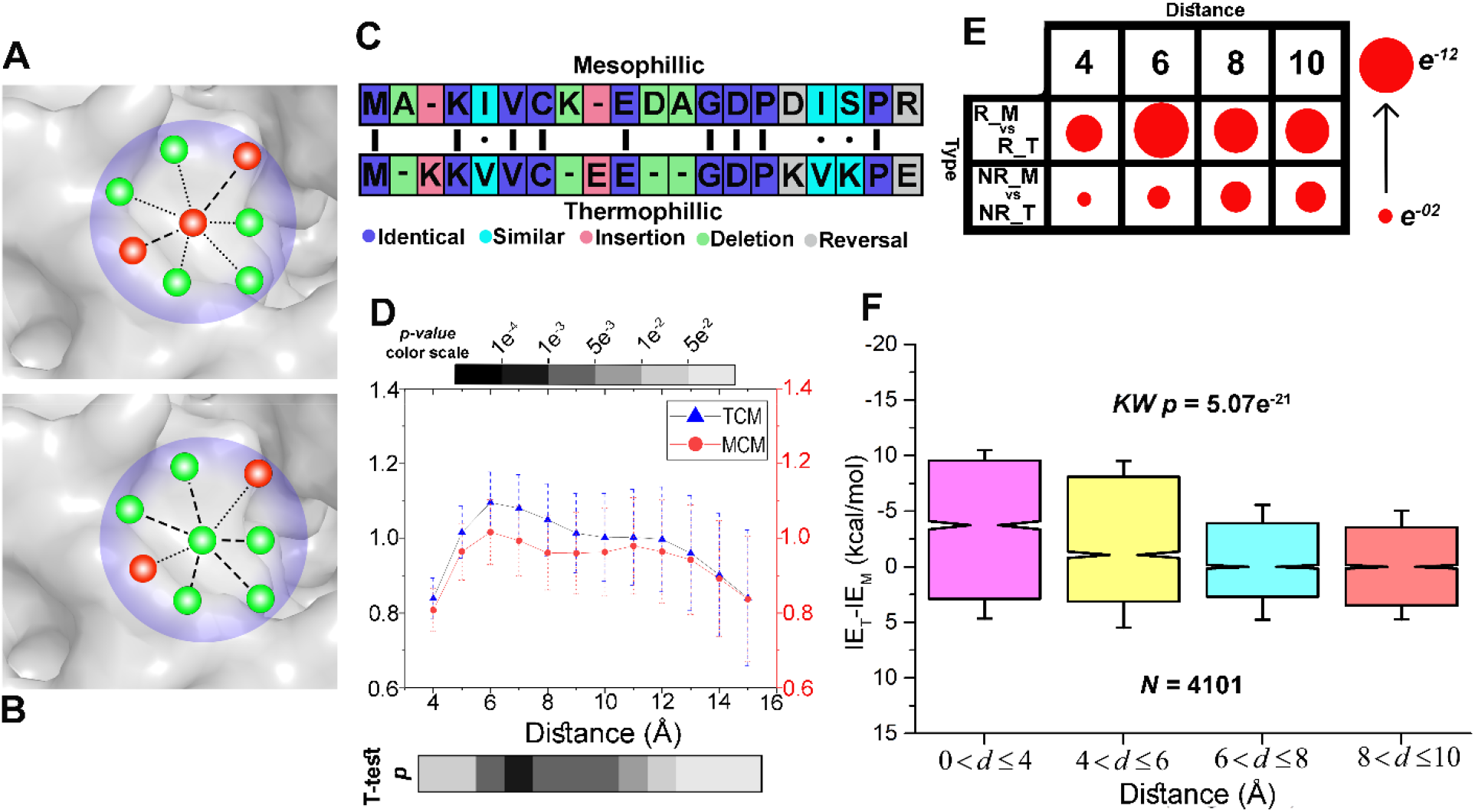
**(A-B)** A schematic diagram showing how a single charge reversal can completely change the existing interaction pattern. Here, the positive and negative charges are represented as red and green spheres respectively. The attractive and the repulsive interactions are represented as dotted and dashed lines respectively. **(C)** It shows how putative charge reversal mutations are found from the pairwise alignment of mesophilic and thermophilic orthologous proteins. **(D)** Comparison of charge-metrics at different distance cut-offs in M-T/HT orthologous proteins. The t-test *p* value for each distance cut-off is represented by a grey scale gradient shown below. **(E)** Interaction energies of charged reversed and non-reversed amino acids in thermophilic proteins have been statistically compared with those in mesophilic proteins using Mann-Whitney U (MW) test. MW *p* values are represented as red circles, where size of the circles represent approximately the level of significance (respective box plots and their *p* values are provided in **Supplementary Fig 2A-D). (F)** Thermo-meso energy difference (*IE*_*T*_ − *IE*_*M*_) associated with the same set of CRMs at different distance bins (*d*) are statistically compared using Kruskal Wallis test.

Charge reversal is a specific type of mutation where charged amino acid in one sequence is substituted by an amino acid with opposite charge in another sequence of a pair-wise sequence alignment (**Fig 1C**). Previous efforts of charge-reversal didn’t find any stabilizing effect. In their study in 1982, Hollecker and Creighton converted all the amino groups to acids groups of a few proteins (14) that can results in the disruption of the salt-bridges, and creation of some new repulsive interactions with the previously present acid groups, hence, found no stabilizing effect. In another study in 2003, Schwehm et. al reversed all the surface charges of a protein and found a destabilizing effect (15), because reversing all the charges at an instance is not a feasible option for gaining thermal stability, as it would disrupt many native interactions and may interfere with the protein’s function. So, in the current study we assess whether the CRMs naturally occur in putative M-T/HT orthologous proteins. If yes, what are the possible roles of CRMs in thermal adaptation? To address this, we have used a carefully curated dataset from our previous study of 1550 M-T/HT naturally occurring putative orthologous protein pairs that (i) share common evolutionary ancestry and exhibit similar (ii) lengths, (iii) domain content and (iv) 3D topologies. Statistical comparison of the effect of putative CRMs further exhibits that charge reversal of a few buried and moderately exposed amino acids may contribute substantially in enhancing stability, which contradicts with the outcome of previous studies (14, 15). Moreover, there are other stabilizing factors for the surface CRMs, like-enhancement of solvent interaction, which weren’t considered in previous studies. This work in its limited scope could be a step forward to understand the broader role of charge reversal in thermostability and provide a new perspective for protein engineers.

## MATERIALS AND METHODS

### Data Collection

Evolutionary expansion of protein families involves several innovations that directly or indirectly affect their domain architectures, functions and global topologies. So statistical comparison of thermophilic and mesophilic putative orthologous proteins to find out thermal adaptation signature often lead to inconsistent results because it is difficult to assess whether the observed differences between orthologs are associated with thermal adaptations or evolutionary innovations. Taking this into account, in our previous study (6), starting from collecting a set of 3346 mesophilic, 988 thermophilic and 407 hyperthermophilic x-ray crystallographic structures (≤ 3.0 Å resolution, and ≥80% protein sequence coverage) from Protein Data Bank (PDB) (16), we constructed a carefully curated datasets of M-T/HT orthologous proteins that not only share common evolutionary ancestry (reciprocal BLAST (17) search with ≥ 50% query sequence coverage and 1^-10^ expected threshold), also exhibit similar lengths (L), 3D topologies and domain architectures. In the current study, we have used a dataset of 1550 M-T/HT orthologous protein pairs (**Data S1**) that exhibit similar length and ACO (18, 19) (ACO, a widely used topological parameter, is calculated as the average separation of contacting amino acids in the primary chain).

### Sequence and structural analysis

#### Pairwise Sequence Alignment of putative orthologs

Pairwise local alignment on 1550 putative orthologous pairs was performed using Water program (20), implemented in EMBOSS Package (21) with a gap-opening and gap-extension penalty of 10.0 and 0.5 respectively. These alignments are used to extract mesophilic-to-thermophilic/hyperthermophilic charge reversal point-mutations within the aligned regions (**Data S1**).

#### Solvent accessible surface area

Solvent accessible surface area (*SASA*) of charged amino acids in the protein structures is computed using Surface Racer tool (22) using a probe radius of 1.4 Å roughly similar to that of a water molecule.

#### Charge-charge and Cation–π interactions

The interaction energy (*G*) for an ion pair is calculated as

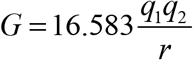

In the above expression, charges (*q*_1_ and *q*_2_) are given in units of electron/proton charge at *pH* 7 (23), distances (*r*) are given in angstroms (Å), and interaction energies (*G*) are obtained in kilocalories per mole (kcal/mol); the numerical factor 16.583 is obtained by the vacuum permittivity, proteins’ average dielectric constant of 20 (24), electron/proton charges in units of coulomb and the conversion factor for energy from joules to kcal/mol. To calculate the total energy, associated with charge-reversal mutations (CRMs) in a protein, all the putative CRMs are first detected from its alignment and then the algebraic sum of *G* s of CRMs with their nearby charges are taken as shown in **Fig 1A-C**.

Cation–π interaction occurs between positively charged side chains of amino acids, Lysine and Arginine and the aromatic rings of Phenylalanine, Tryptophan and Tyrosine (that are placed approximately 4 Å apart), is believed to be a key contributor to proteins folding and thermal stability (25). Our own in-house python scripts are used to identify these interactions.

### Statistical analysis and Figure Preparation

PAST (PAleontological STatistics) software (26) and our own in-house python scripts are used for the statistical analyses. Test used, number of data points and respective *p*-values are provided in the plots. All the images are produced using OriginPro and Seaborn package (Michael Waskom and the Seaborn development team) of Python 2.7.

## RESULTS AND DISCUSSION

In order to function in the extreme environments, thermophilic proteins have been designed differently than mesophilic proteins (1). This designing principle includes changes in the sequence level that is further associated with several secondary and tertiary structural factors of thermostability (6). Thermophilic proteins feature a higher number of charge-charge interactions than their mesophilic orthologs (6, 27–30), which may be attributed to the fact that thermophilic sequences are generally more enriched with charged residues (6, 31, 32). Moreover, each amino acid can participate in several attractive and repulsive interactions simultaneously. Henceforth, the (attractive – repulsive) interactions per charged residue (charge metric (33), *CM*) may provide a more appropriate signature than counting charge-charge interactions. Here, we have calculated the *CM* for thermo (*TCM*) and meso (*MCM*) for varying distance cut-offs from 4 Å to 15 Å, and we observe that *TCM* and *MCM* significantly differ up to a distance of 12 Å (Two sample t-test *p* value ≤ 0.01) (**Fig 1D**). Since interaction energy of any charge amino acid pair at 11 and 12 Å becomes negligibly smaller (**Supplementary Fig 1**), distance cut-offs up to 10 Å are considered for further study.

### CRMs possess more stabilizing energy in thermophilic proteins at short range

We started with extracting all the meso to thermo charge-reversal point mutations for all 1550 orthologous pairs from their alignments (**Fig 1C**). Net interaction energy (*IE*, where *IE* = Σ*G*) for amino acids in mesophiles are distributed in two groups: R_M for reversed and NR_M for not reversed, similarly, R_T and NR_T for thermophiles. Mann Whitney U (MW) test is performed for R_M, R_T and NR_M, NR_T for all distance cutoffs from 4 Å to 10 Å. At every cutoff, the MW *p* value of comparison in R_M and R_T is more significant than that of NR_M and NR_T (**Fig 1E, Supplementary Fig 2A-D**). Thermophilic proteins are enriched with charged residues (31, 32) whose positions in the 3D structure are such that in turns they create more stabilizing charge-charge interactions (27–30) than mesophilic proteins. Among all the charged residues in thermo, putative CRMs contribute even more in stabilizing energy gain than the rest, and this trend is stronger at short distance cutoffs **(Fig 1E, Supplementary Fig 2A-D)**.

A comparison of *IEs* of the common CRMs in different distance bins of 0-4, 4-6, 6-8, 8-10 Å shows that the average interaction energy gain (*IE*_*T*_ − *IE*_*M*_) per CRM (−1.54 kcal/mol, -1.36 kcal/mol, -0.39 kcal/mol, and -0.14 kcal/mol at the bins respectively) significantly differ from one another (Kruskal Wallis, (KW) *p* value = 5.07*e*^−21^) (**Fig 1F**). This indicates that CRMs do not contribute equally at every distance and the overall energy gain arises mainly from the interactions at a shorter distance (≤ 6 Å). Beyond that distance the attractive and repulsive interaction energies associated to the CRMs nearly counter balance each other in both thermophilic and mesophilic proteins. Since, more than one CRM may occur in a meso-thermo orthologous protein pair, how they collectively contribute in the stability of the protein will give more realistic picture of their role in thermostability.

To understand how the CRMs contribute in thermostability at protein level we estimated two parameters for each meso-thermo orthologous pair, 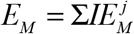 and 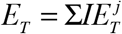 that represent the total interaction energy for meso and thermo, where 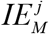 and 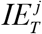 stand for the energy associated with *j*^*th*^ CRM in meso and thermo respectively. In these comparisons, our null hypothesis *H* _0_ is *E*_*T*_ = *E*_*M*_ and the alternative hypothesis *H*_1_ is *E*_*T*_ > *E*_*M*_ for the statistical test at each distance bins. The difference is more significant for the 0-4 Å (MW *p* value =1.44*e*^−73^) and 4-6 Å (MW *p* value = 3.06*e*^−71^) bins compared to 6-8 Å (MW *p* value = 4.32*e*^−52^) and 8-10 Å (MW *p* value = 7.04*e*^−15^) bins (**Fig 2A, Supplementary Fig 3A-D**). This result indicates that the contribution of CRMs is limited to a larger extent on the short-range interaction compared to long-range interactions. The difference of *E*_*T*_ and *E*_*M*_ (*E*_*T*_ − *E*_*M*_) plotted for the distance bins and their statistical comparison (Kruskal Wallis (KW) *p* value = 3.62*e*^−74^) further supports the previous result (**Fig 2B**).

**Fig 2.**
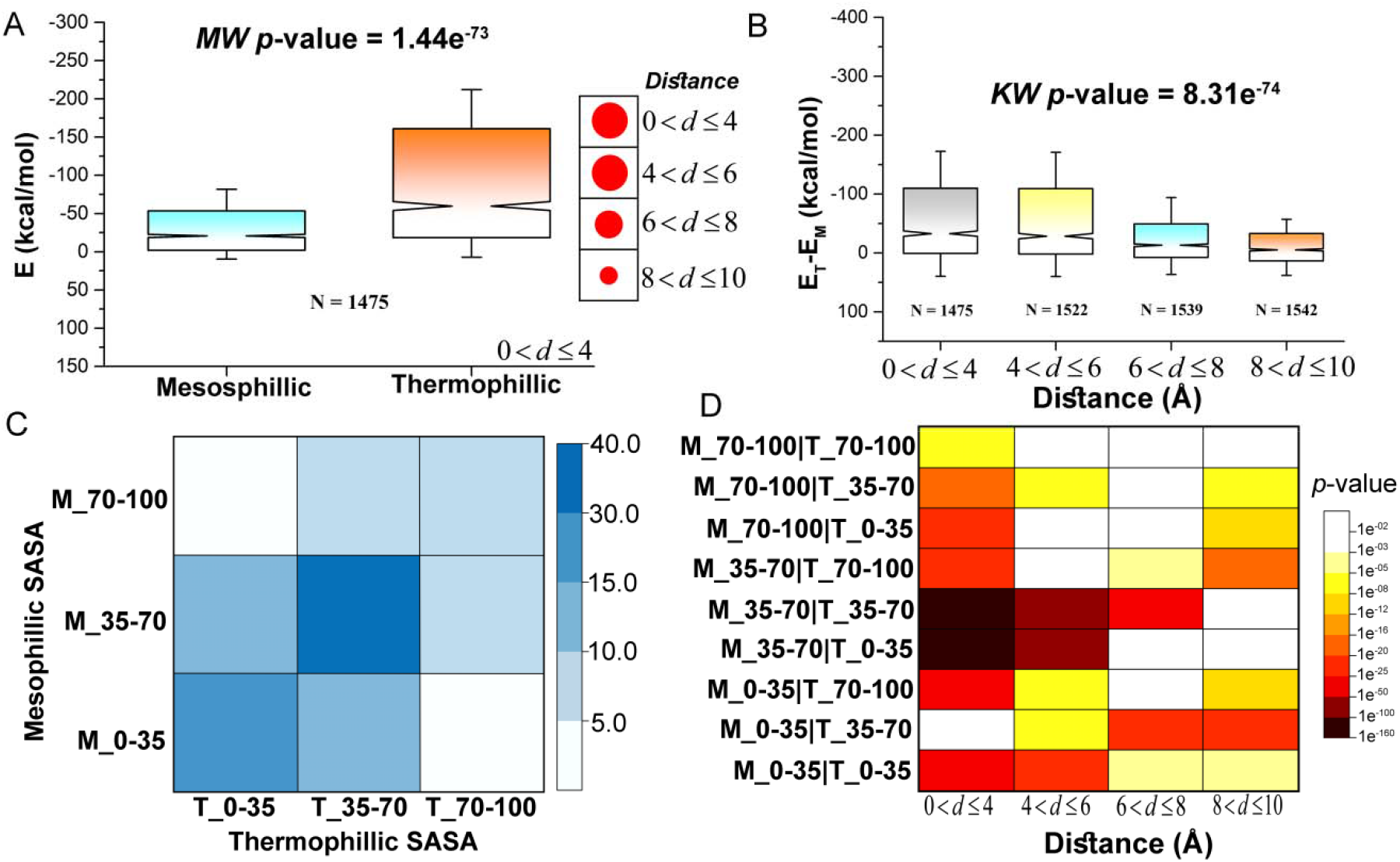
**(A)** Statistical comparison (MW test) and its p value of total energy associated with CRMs in mesophilic proteins and their respective thermophilic orthologs for the distance bin 0-4 Å is provided. For the rest of the distance bins the *p* values (represented as red circles, where there sizes roughly represent the significance level) of MW test are provided alongside. Box plots for all the distance bins are given in **Supplementary Fig 3A-D. (B)** Stabilizing energy gain associated with CRMs for the M-T/HT orthologous proteins at different distance bins are compared using Kruskal Wallis test. **(C)** Occurrence of CRMs in different Solvent Accessible Surface Area (SASA) ranges in M-T/HT orthologous proteins are represented as heatmap. A darker shade of blue represents higher percentage of occurrence. **(D)** MW *p* values of comparisons of interaction energies associated to CRMs for M-T/HT orthologous pairs at different SASA ranges at different distance bins are represented as heatmap. Level of significance increases from white to dark-brown.

### Majority of the total stabilizing energy gain appears from the short range interaction by buried and moderately exposed CRMs

Charged amino acids prefer to occur near the surface in both mesophilic and thermophilic. However, this tendency is stronger in thermophiles as it contributes by increasing solvent interactions (34–36). On the other hand, burial of charged residue comes with desolvation entropic penalty which may be reduced in thermophilic by proper networking of charged residue (37). To investigate where in the structure the CRMs occur, we distributed all the CRMs in three groups based on their solvent accessibility i) buried (Charged solvent accessible surface area, C_SASA ≤ 35%, M_0-35 and T_0-35), ii) moderately exposed (35% < C_SASA ≤ 70%, M_35-70 and T_35-70), and iii) exposed (70% < C_SASA ≤ 100%, M_70-100 and T_70-100), where M and T stand for meso and thermo respectively. Number of mutations occur in the moderately exposed group account for ∼55% of the total mutation in both meso and thermo (**Fig 2C**). However, CRMs may fall in other groups in meso and thermo. Still the amino acids that remain moderately exposed in both (36.2%) account for more than two fold than the next populated group, where they are buried in both (16.6%) (**Fig 2C**). Total energy comparison of these sets of mutations for orthologous meso-thermo pairs, using the same methodology described above, demonstrates that optimization of the short-range interactions occurs predominantly at the moderately exposed and buried segments of the protein structure (**Fig 2D**). Here, we hypothesise that rational optimization of moderately exposed short range electrostatic interaction by charge-reversal may be used for engineering thermostable variant of mesophilic proteins.

### Additional factors

However, only 75.3%, 72.5%, 66.3% and 57.6% of the total orthologous pairs exhibit energy gain associated with CRMs at 0-4, 4-6, 6-8, and 8-10 Å bins (**Supplementary Fig 4**) respectively. For those who possess energy loss we looked for other counter balancing factors. Here, we limit our discussion to the effect of CRMs only. Comparison of cation-pi contacts for energy loss associated M-T/HT orthologs exhibit no significance difference. Another possibility is the interaction of the charged amino acids with its solvent environment. Exposure of charged groups on protein surface facilitates thermal stability by blocking penetration of the solvent within the protein. This is achieved by a highly connected network of water-water hydrogen bonds, attached to the protein surface by amino acids-water hydrogen bonds (34– 36). Irrespective of energy gain or loss cumulative solvent accessibility of charge residues in thermophilic proteins exhibit more C_SASA compared to those in their mesophilic orthologs (**Fig 3A**). However, when considered individually, energetically destabilizing CRMs (i.e. those associated with interaction energy loss) in thermophilic proteins exhibit more exposed surface than stabilizing CRMs, indicating a counter balancing role of C_SASA locally (**Fig 3B**).

**Fig 3.**
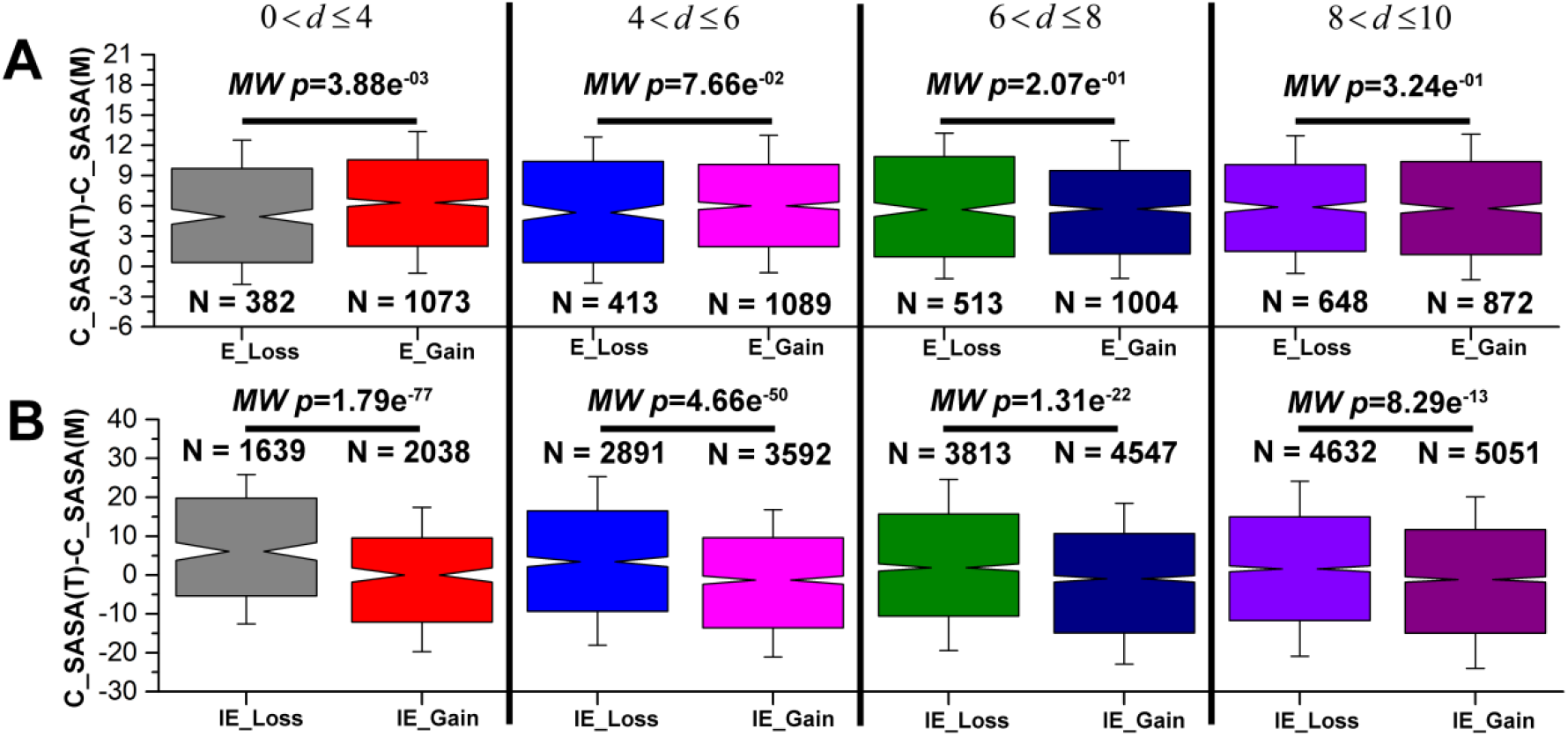
**(A)** Difference of the percentage of SASA occupied by charged amino acids in thermophilic and mesophilic orthologous proteins are plotted for the pairs that exhibit a gain and loss in interaction energy due to CRMs at different distance bins. MW test is performed for each set and their *p* values are provided. **(B)** Difference of the percentage of area of CRMs, exposed to the surface in thermo and meso are compared where a gain and loss in interaction energy is found due to CRMs at different distance bins. MW test is performed for each set and their *p* values are provided.

In our study, working on a large, carefully curated dataset, we aim to understand, in general, how CRMs contribute in thermostability. However, when we calculate the charge-charge interaction energy, all the interactions are treated equally, although several molecular dynamics and experimental studies show that the interaction energy depends on several other factors like location of the charges on the ions, positions of the amino acids in the structure, dielectric constant, folded and denatured state of the protein etc (33, 38, 39). Moreover, dielectric constant varies from core to surface and also with temperature (40). However the inclusion of the effect of the variation of dielectric constant and temperature will not alter the conclusion of our study.

## Conclusion

While strategies for attaining thermal stability and its application in protein engineering have been studied for decades, CRMs remain unpracticed. In our study, statistical comparison of sequence and structural features and semi-empirical energetics of 1550 M-T/HT orthologous protein pairs with remarkable structural and topological similarity reveals the following key points that could be useful for protein engineers as a future reference.

1. CRMs prevalently occur in naturally occurring putative orthologous pairs. Not only that, they contribute subsequently to protein stability by altering short distance charge-charge interactions. Hence, CRM acts as one of the key mechanisms for gaining thermal stability.
2. Positional comparison of interaction energy of the CRMs exhibits higher energy gain for buried and partially exposed CRMs compared to those who are exposed to the solvent environment.
3. CRMs associated with energy loss are more solvent-exposed than the rests, indicating a higher role of solvent interaction in the former.

## Acknowledgement

The authors would like to acknowledge Raktim Maity and Soumya Kundu for their valuable comments during the discussions.

## Author Contribution

S.H. and S.K. designed research; S.H. implemented computational methodologies, S.H. and S.B. performed research; S.H., S.B. and S.K. analyzed data and wrote the paper.

