## Supplementary Figures for "Can charge-reversal be considered as a strategy for attaining thermal stability in proteins?"

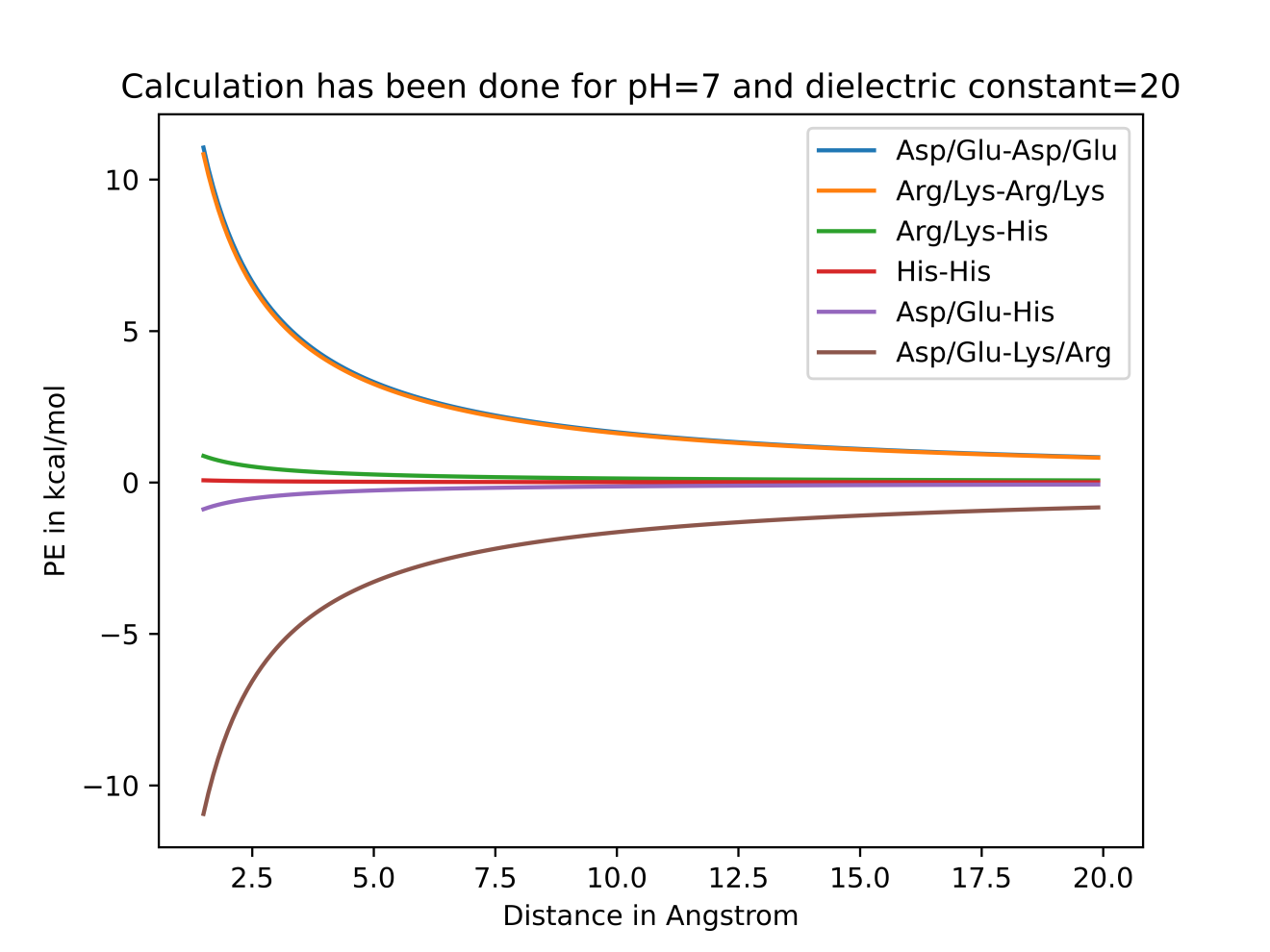

**Figure S1.** The potential energies of charge-charge interactions for each pair of charged amino acids for pH=7 and dielectric constant of 20 at different distances (in angstrom).

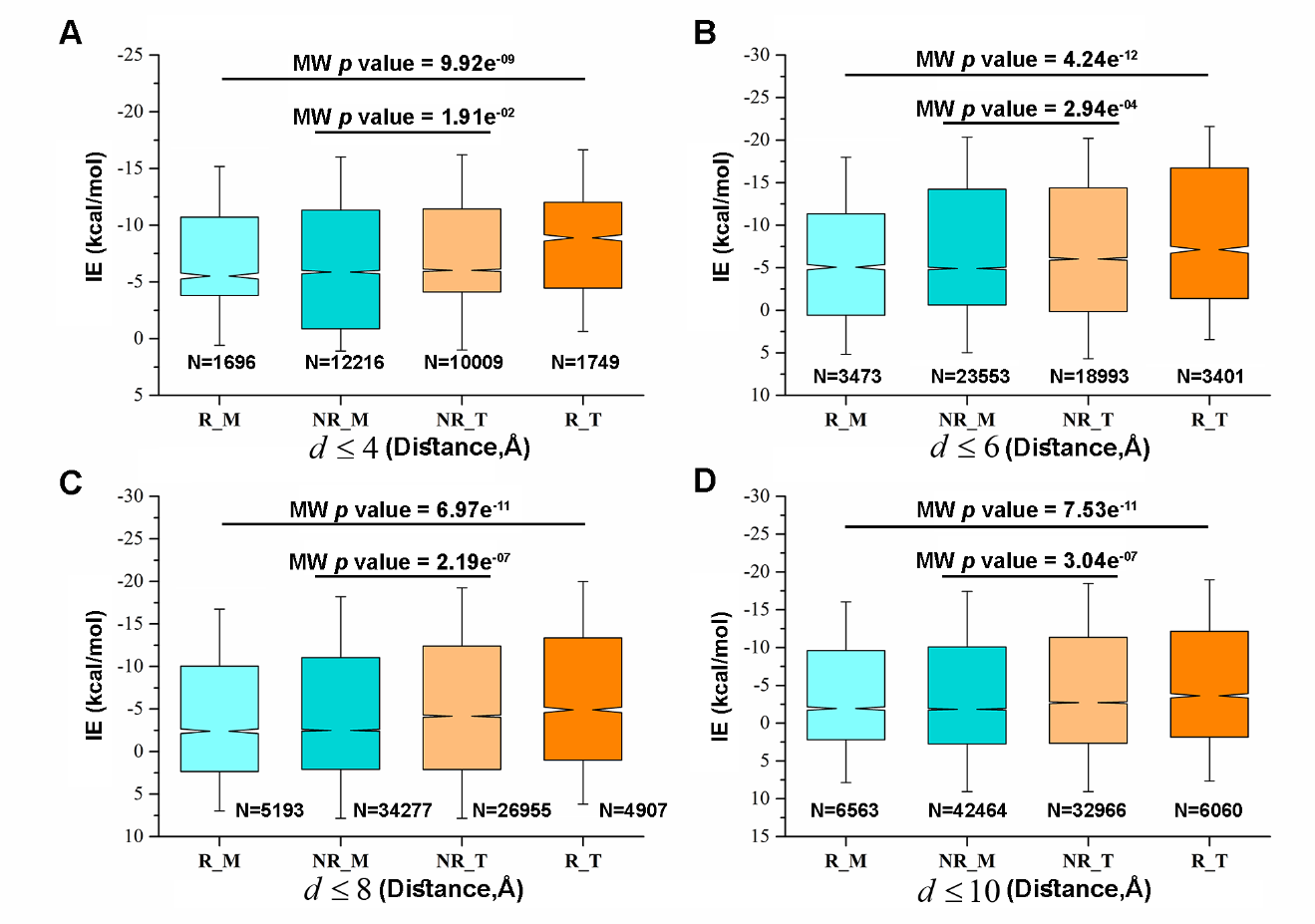

**Figure S2.** Charged amino acids in each of thermo and meso are distributed in two groups charged reversed (R_T and R_M respectively) and not reversed (NR_T and NR_M respectively). The distribution of electrostatic interaction energy of charged amino acids (IE in kcal/mol) for different distance (*d*) cutoffs are represented as box plots. Mann-Whitney U test *p* values are provided for comparisons of R_T, R_M and NR_T, NR_M for every distance cutoffs.

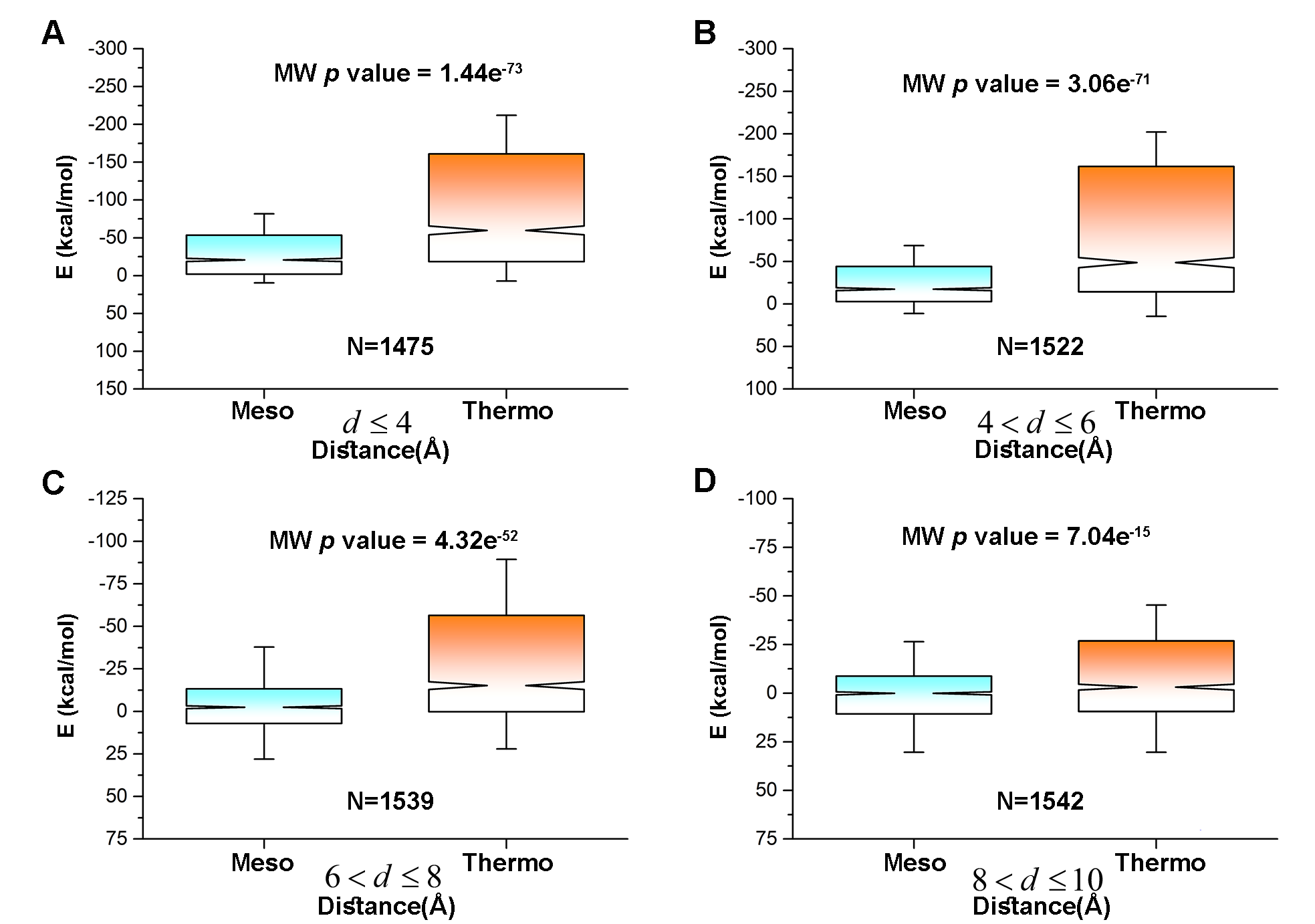

**Figure S3.** The distribution of electrostatic interaction energy of Meso-Thermo/Hyperthermo (M-T/HT) orthologous proteins associated with charged reversed amino acids (E in kcal/mol) for different distance bins are represented as box plots. Mann whitney U test *p* values are provided for comparisons of energy distribution of (M-T/HT) orthologous protein pairs for different distance bins.

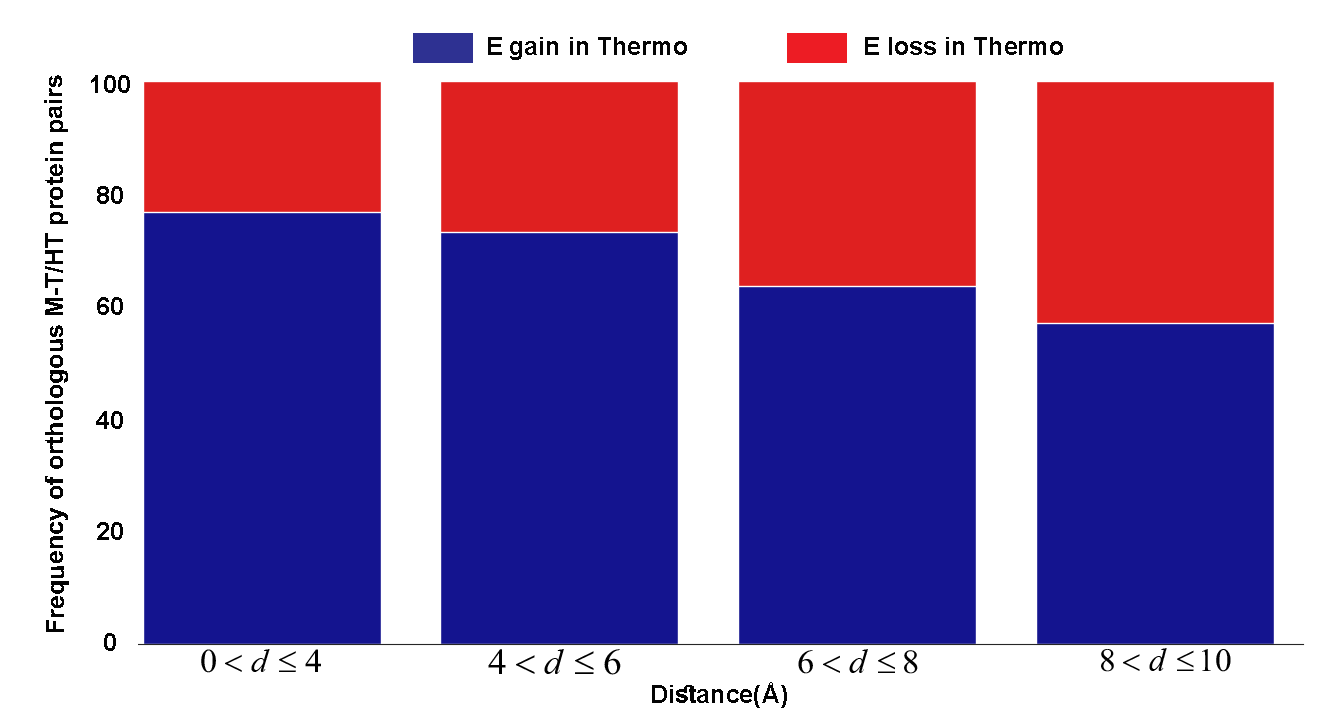

**Figure S4.** Frequency M-T/HT orthologous protein pairs that exhibit interaction energy gain or loss due to charge reversal are provided for different distance bins.
